# Nonlinear progression during the occult transition establishes cancer lethality

**DOI:** 10.1101/2024.04.23.590826

**Authors:** Joshua D. Ginzel, Henry Chapman, Joelle E. Sills, Edwin J. Allen, Lawrence S. Barak, Robert D. Cardiff, Alexander D. Borowsky, H. Kim Lyerly, Bruce W. Rogers, Joshua C. Snyder

## Abstract

Cancer screening is based upon a linear model of neoplastic growth and malignant progression. Yet, historical observations suggest that malignant progression is uncoupled from growth which may explain the paradoxical increase in early-stage breast cancer detection without a dramatic reduction in metastatic burden. Here we lineage trace millions of genetically transformed field cells and thousands of screen detectable and symptomatic tumors using a cancer rainbow mouse model of HER2+ breast cancer. Transition rates from field cell to screen detectable tumor and then to symptomatic tumors were estimated from a dynamical model of tumor development. Field cells are orders of magnitude less likely to transition to a screen detectable tumor than the subsequent transition of a screen detectable tumor to a symptomatic tumor. Our model supports a critical occult transition in tumor development during which time a transformed cell becomes a *bona fide* neoplasm. Lineage tracing and test-by-transplantation reveals that nonlinear progression during or prior to the occult transition gives rise to nascent lethal cancers at screen detection. Simulations illustrate how occult transition rates are a critical determinant of tumor growth and malignancy in the lifetime of a host. Our data provides direct experimental evidence that cancers can deviate from the predictable linear progression model foundational to current screening paradigms

## Introduction

Progression of a screen detectable early-stage tumor to a symptomatic and metastatic tumor is often regarded as the pivotal moment in the natural history of cancer^1^. As such, most cancer screens are focused on identifying and treating small tumors prior to their growth and progression to metastasis. Although this concept has led to remarkable progress in cancer screening programs, the cost-benefits of screening are being questioned^2^. For instance, not all screen-detectable ductal carcinomas (DCIS) of the breast progress to invasive ductal carcinoma (IDC) and despite significant progress in early detection, metastatic burden remains relatively unchanged^2,3^.

Leslie Foulds proposed six-rules of tumor progression more than seventy years ago. Foulds’ rules conceptualized that progression to malignancy occurs independent of neoplastic growth, can occur by multiple developmental trajectories, and can occur abruptly or gradually. Thus, it should not be surprising how screening for early-stage tumors by size alone may not produce optimal benefits. Yet, the absence of quantitative proof of Foulds’ rules in immune intact mammalian models continues to confound our understanding of critical inflection points in the disease process. One major challenge is that most of the disease process occurs over the course of several years in humans and is dependent upon cancer cell extrinsic adaptations in the tumor microenvironment^4,5^. Tumor driver gene mutations also occur several years prior to detection and many fully transformed clones can stall or extinguish entirely without ever causing disease^6–9^. As a result, for every transformed cell that develops into cancer, billions of transformed cells exist in healthy tissues that may never form a tumor in a patient’s lifetime^10^. Altogether, this makes solving for the tumorigenic potential of a transformed cell and its progression to a metastatic cancer extremely challenging in a complex biological organism.

We previously developed Cancer rainbow mouse (Crainbow) models for fluorescently visualizing and barcoding premalignant clones^11,12^. Our previous work in the mammary gland validated a HER2 Crainbow model that expresses multiple clinically relevant isoforms of the protooncogene HER2 including wild-type (WT), an exon16-splice isoform (d16), and an N-terminally truncated isoform (p95)^12^. We qualitatively found that the clinically apparent and symptomatic phase of tumorigenesis in HER2 Crainbow (HBOW) mice is characterized by isoform dependent differences in tumor phenotypes, suggesting that even in mouse models an early inflection point in the disease process must occur prior to detection. However, to quantitatively understand tumor development and progression requires hundreds to thousands of samples across the entire course of the disease.

Our goal here is to lineage trace and mathematically model tumor growth in order to answer several basic questions that remain unresolved. How many genetically transformed cells in a field are required to form a tumor, how frequently do transformed cells establish screen detectable tumors, and how frequently do these screen detectable tumors transition to clinically apparent and symptomatic tumors? Lastly, we seek to determine whether progression occurs independent of these growth phases. We demonstrate that an occult transition from a genetically transformed cell to a successful tumor is exceptionally rare and is the rate limiting step in the growth of a tumor. Our data show also show that progression is indeed independent of growth and further illustrates how an abrupt progression to malignancy during the occult transition establishes nascent lethal cancers prior to detection.

## Results

Tumorigenesis begins at cell transformation when a tumor driver gene is first expressed. In our model, initiation of HER2 expression occurs no later than two-weeks of age (Figure 1A). Our goal is to develop a model of tumor development that includes growth of transformed cells, the transition to screen detectable neoplasms, and then to clinically detectable symptomatic tumors. Therefore, we first conducted a longitudinal study to determine the duration of the preclinical phase and symptomatic phase of tumorigenesis in HBOW mice. We used palpation as a proxy for clinical detection and surveyed HBOW mice for the first sign of a tumor. Then tumors were measured at least two times weekly until total tumor burden reached 2000mm^3^ or a humane endpoint. At endpoint, mice were imaged to retrieve the fluorescent protein (FP) barcode for each palpable tumor (Figure 1B). Tumor free survival and overall survival were both calculated in twenty-eight mice. HBOW mice have a fifty-percent tumor free survival at 113 days (∼16 weeks of age) and all mice develop tumors by 148 days (∼21 weeks of age) (Figure 1C). Fifty percent of HBOW mice reach tumor burden by 168 days (∼24 weeks of age) and all mice reach tumor burden by 198 days (∼28 weeks of age) (Figure 1D). Thus, the preclinical phase of tumorigenesis as measured from the time of oncogene expression to the first symptomatic tumor is approximately 14 weeks while the clinical or symptomatic phase of tumorigenesis is approximately 55 days (7.8 weeks) demonstrating that over two thirds of the tumor growth phase is missed by palpation only.

**Figure 1.**
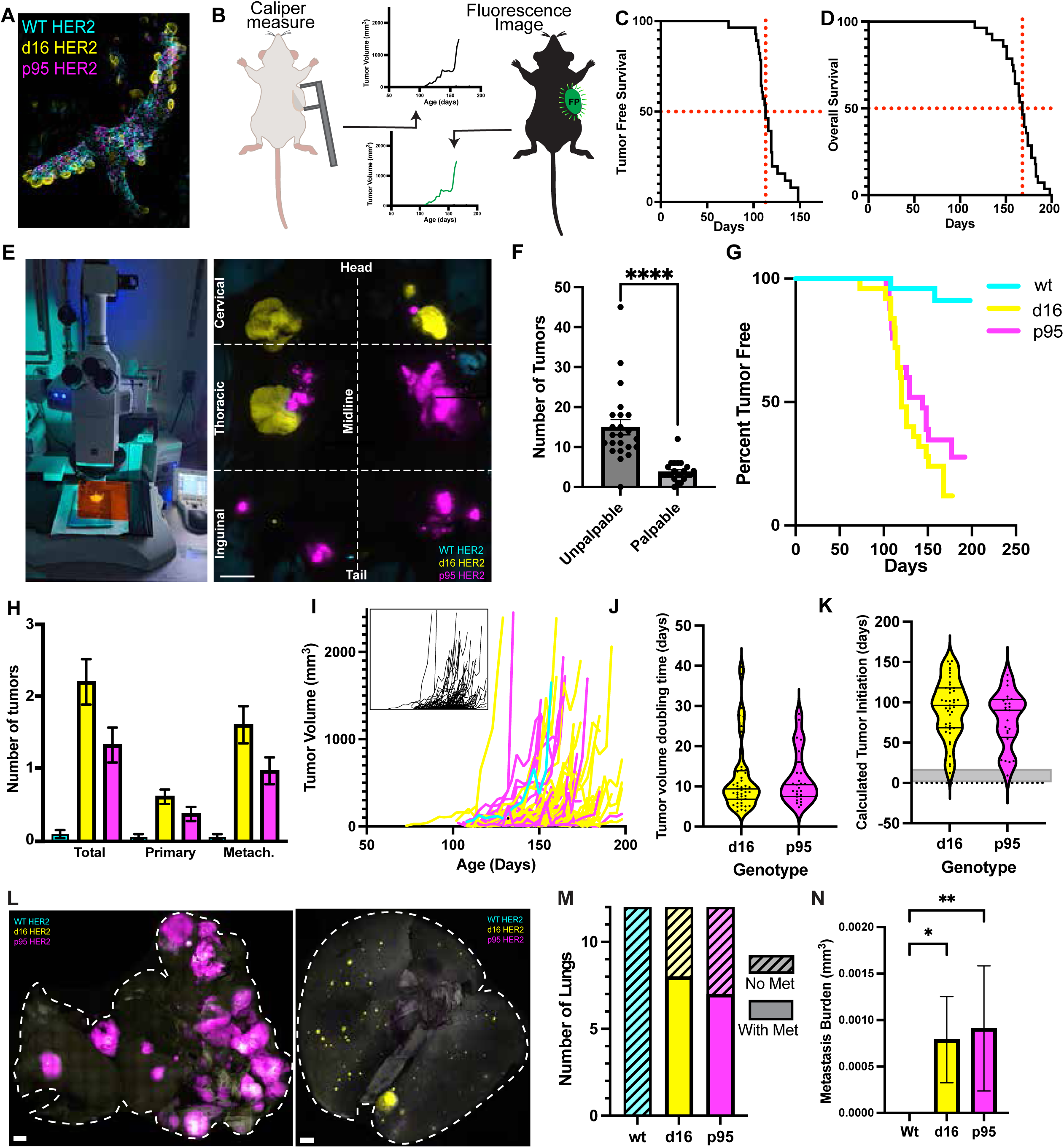
Quantifying the visible phase of tumorigenesis. **A,** Whole mammary gland imaging of fluorescent Crainbow cells expressing mTFP:WT HER2 (cyan), eYFP:d16 HER2 (yellow), mKO:p95 HER2 (magenta) at 2 weeks. **B,** Tumors palpable through the skin were measured twice weekly with calipers until tumor burden endpoint. Mice were imaged at necropsy to retrieve fluorescent protein expression and registered to the palpation data. **C,** Tumor free survival of 28 mice palpated biweekly. Red lines represent 50% on the y-axis and 113 days on the x-axis. **D,** Survival until tumor burden endpoint of 28 mice. Red lines represent 50% on the y-axis and 168 days on the x-axis. **E,** Whole mouse imaging reveals tumor genotype by fluorescence at necropsy. **F,** Number of tumors per mouse measured by whole mouse imaging that were palpable or too small to palpate (unpalpable). Statistical significance was determined by Mann-Whitney test. P value is represented as follows: ****=p<0.0001 **G,** Tumor free survival calculated by the time to the first palpable tumor per mouse of each color as determined by whole mouse imaging. **H,** Number of tumors expressing each HER2 isoform per mouse at tumor burden endpoint (N=28 mice). From left to right: Total tumors observed, Primary tumor observed, All metachronous tumors observed. **I,** Observed tumor volume growth over time colored by genotype. Inset shows growth before Crainbow registration. **J,** Tumor volume doubling(TVDT) time of each observed tumor by genotype. Tumors with <3 measurements were not included. **K,** Calculated time of tumor initiation from tumor volume doubling time and first palpation observation. Gray shade indicates actual recombination and oncogene expression between 0 and 14 days old. **L,** Representative 3D whole lung imaging for HBOW fluorescence. Lung volume outlined with white dotted line. **M,** Number of lungs with at least one metastatic lesion expressing mTFP:WT HER2 (cyan), eYFP:d16 HER2 (yellow), mKO:p95 HER2 (magenta) from mice at tumor endpoint. **N,** Average area of metastatic burden per lung for each genotype. Statistical significance was determined by Kruskal-Wallis test. P values are represented as follows: *=p<0.05, **=p<0.01. Data are represented as Mean +/- SEM. Scale bar = 1mm.

All mice were imaged using an epifluorescent dissecting microscope and palpable tumors were co-registered with tumor genotype using the Crainbow fluorescent barcode (Figure 1E). This allowed us to also determine if differences in the length of the preclinical phase or symptomatic phase could be detected between each HER2 isoform. Whole mouse imaging allows for the detection of a significant reservoir of preclinical lesions, undetectable by palpation, that were approximately 3-fold more abundant than palpable lesions and as small as 0.5mm in diameter (Figure 1F). Of the ninety palpable tumors, there were two WT expressing tumors (cyan:TFP1), fifty-five d16 expressing tumors (yellow:EYFP), and thirty-three p95 expressing tumors ( magenta:mKO). Using this registration, tumor free survival was plotted for each HER2 isoform. Twenty of the surveyed mice had at least one palpable d16 tumor with a median tumor free survival of 120 days (∼17 weeks) compared to the 17 mice with at least one palpable p95 tumor and an increased median time of 144 days (∼20 weeks) (Figure 1G). On average there were 2.2 d16 tumors, 1.32 p95 tumors, and 0.08 WT tumors per mouse (Figure 1H). HER2 Crainbow mice develop multifocal tumors, so we also reported on the lineage of first palpable tumors and compared this to the lineage of metachronous palpable tumors. First palpable tumors (primary) accounted for twenty-seven percent of total tumor burden, whereas the remaining seventy-three percent were metachronous tumors that developed independent of the first tumor. Similar frequencies for tumor penetrance for each genotype were observed for primary palpated tumors and metachronous tumors (Figure 1H). Altogether, this data demonstrates that the length of the preclinical phase is similar between d16 and p95 and also confirms that WT HER2 has limited tumorigenic potential.

Next, growth rates were calculated for d16 and p95 tumors. WT was not included due to the limited number of WT tumors. For each tumor palpated over time, measurements were plotted and registered to the genotype (Figure 1I). Tumor volume doubling time(TVDT) is a commonly used tool to asses tumor growth rate and has been correlated to tumor phenotype ^13,14^. TVDT was on average 12 days for both p95 HER2 and d16 HER2 tumors (Figure 1J). Assuming that the TVDT is constant, the age of tumor initiation can be calculated by using the first palpable measurement and the calculated TVDT. Inferred tumor initiation is calculated to be 93 days (13 weeks) for d16 tumors and 78 days (11 weeks) for p95 tumors (Figure 1K). However, in our system, oncogene initiation and cell transformation occurs no later than 14 days of age. Thus, palpation and TVDT alone is a poor predictor of tumor initiation and an inadequate method to measure tumor development. Similar to humans the majority of the disease process occurs prior to clinical detection and as such cannot be estimated from growth during the clinical phase.

Metastatic potential was also measured by 3D whole lung imaging at tumor burden endpoint (n=12). Representative images show high p95 and d16 metastatic burden in the lungs of two different mice (Figure 1L). Of the 12 lungs, none had WT metastasis, eight had d16 metastases, and seven had p95 metastases (Figure 1M). The average area of metastatic lesions from whole lung images revealed that the metastatic burden for d16 and p95 were similar (Figure 1N) and demonstrated that each isoform has the potential to drive tumors that progress to metastasis.

The clinical or symptomatic phase appears similar for d16 and p95 tumors; each tumor genotype grows at similar rates and is capable of progression to a metastatic cancer. However, major differences in the preclinical phase have been overlooked and preclude a quantitative assessment of the actual tumorigenic potential of a single HER2 transformed cell. Simply stated, do d16 and p95 transformed cells grow and progress similarly? We reasoned we would need to know the number of initial cells transformed, their growth as fields over time, and the rate of production for early screen detectable tumors (unpalpable) and symptomatic tumors (palpable).

First, we began by collecting data on the number of transformed cells at initiation, their growth, and the overall carrying capacity of transformed cells in the mammary gland prior to screen detection. Quantitative imaging and analysis were performed in HBOW mice sacrificed at 3, 4, 6, and 10 weeks of age. Single cell suspensions from every gland (10 glands / mouse) were prepared and a representative sample of cells counted by confocal to estimate total cells by HER2 isoform. The total carrying capacity – that is the number of epithelial cells present within the duct – increased by almost three-fold from 3 weeks to adulthood as expected due to the pubertal development of the mammary duct (Figure 2B). Concomitant increases in WT and d16 cell numbers were observed through 6 weeks but p95 cell numbers remained stagnant and only started to variably increase at 10 weeks of age due to the fact that some mice start to develop very small screen detectable lesions at this age (Figure 2B,C). Throughout gland development, d16 HER2 expressing cells make up roughly half of the recombined mammary epithelium while WT HER2 increases from 30% to 44% during pubertal development (Figure 2C). In contrast, p95 HER2 expressing cells fall from 22% to 7% of the recombined epithelium during puberty and variably increase to 14% on average at 10 weeks of age (Figure 2C).

**Figure 2.**
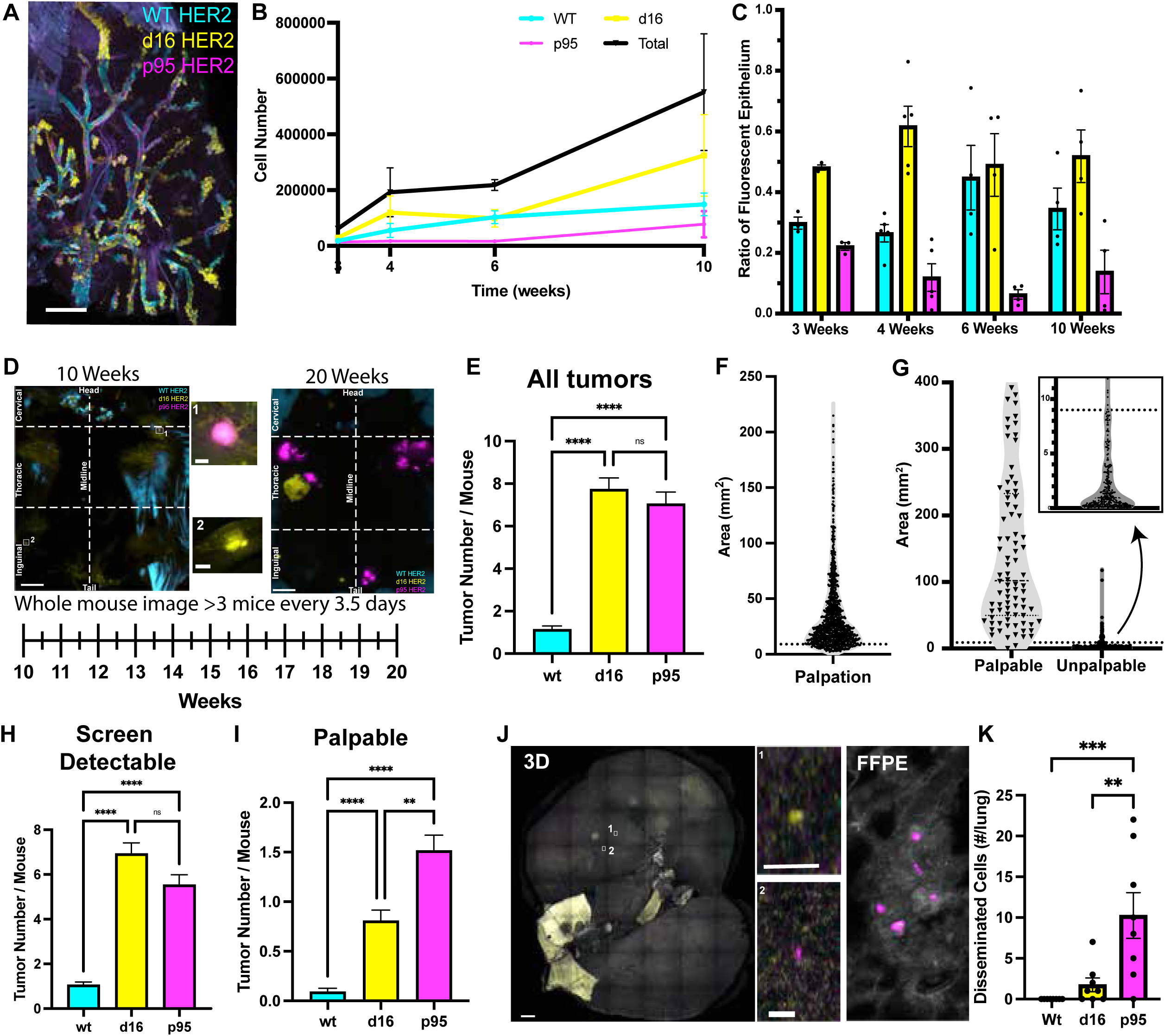
Quantifying the invisible phase of tumorigenesis. **A,** Whole mammary gland imaging of fluorescent Crainbow cells expressing WT (cyan), d16 (yellow), p95 (magenta) at 10 weeks of age. **B,** Quantification of the number of fluorescent cells from all ten mammary glands of HBOW mice after dissociation to single cells at 3, 4, 6, and 10 weeks of age (n=2-3 mice). **C,** Proportion of cell number for each genotype from **B.D,** Whole mouse imaging of HBOW mice at 10 weeks of age and 20 weeks of age shows representative tumor number and size. To survey occult tumor growth, HBOW were sacrificed every 3.5 day from 10 weeks to 20 weeks of age to survey tumor size and genotype. **1&2** are high magnification insets from the 10 week old whole mouse image. **E,** Quantification by genotype of all tumors based on Crainbow fluorescence from whole mouse imaging (n= 120 mice). Statistical significance was determined by Kruskal-Wallis test. **F,** Calculated area of each palpation measurement taken in Figure 1C,D. Dotted line marks the 15^th^ percentile at 9mm^2^. **G,** Tumor area from whole mouse imaging of palpated mice in Figure 1D,E. Dotted line marks the putative cutoff between palpable and unpalpable at 9mm^2^. Inset shows the number of unpalpable lesions that fall below the 9mm^2^ cutoff. **H&I,** Quantification by genotype of screen detectable tumors less than 9mm^2^ **(H)** and palpable tumors greater than 9mm^2^ (**I**) based on Crainbow fluorescence from whole mouse imaging (n= 120 mice). Statistical significance was determined by Kruskal-Wallis test. **J,** Representative image from whole lung imaging of HBOW mice at 10-12 weeks of age. Inset: High magnification imaging shows single disseminated cells in the lung expressing p95 (magenta) or d16 (yellow). **K,** Quantification of the number of disseminated cells from whole lung imaging at 10-12 weeks of age (n=8) Statistical significance was determined by One-way ANOVA with Bonferroni’s multiple comparison test. **E,G,J&K:** **=p<0.01, *** = p<0.001, **** = p<0.0001. Data are represented as Mean+/- SEM. Scale bar = 1mm for **A,D,F**; 0.4mm for **F insets**.

Next, we quantified the number of screen-detectable tumors by examining the total number of small, unpalpable lesions over time that were only detectable by whole mouse imaging. HBOW mice were necropsied and imaged at time points every 3-4 days from 10 to 20 weeks of age to quantify unpalpable, image detectable lesions immediately after the carrying capacity of the field is reached (10 weeks) and up to the median time of clinical detection (20 weeks) (Figure 2D). Representative images at 10 weeks show a few small lesions within the Crainbow labeled mammary ducts (Figure 2D insets 1-2). These lesions continued to increase in number and size through 20 weeks (Figure 2D). Image detectable tumor burden for d16 HER2 and p95 HER2 was nearly identical, and each were seven-fold more likely than WT HER2 (Figure 2E). In order to differentiate between an early palpable lesion and an unpalpable one from whole mouse imaging alone, volume measurements from the previous palpation experiments were converted to area and used to find the lower 15^th^ percentile of palpable tumors (9mm^2^)(Figure 2F). Applying this cutoff to whole mouse imaging from Figure 1E&F revealed that 98% of palpable tumors and 88% of unpalpable tumors were correctly classified (Figure 2G). After stratification of whole mouse imaging data using the 9mm^2^ cutoff, mice had on average 1 WT tumor, 7 d16 tumors and 5.5 p95 HER2 tumors that were smaller than 9mm^2^ (Screen detectable, Unpalpable:S) and 0.09 WT tumor, 0.8 d16 HER2 and 1.5 p95 HER2 tumors that were larger than 9mm^2^ (Large, Palpable:L) (Figure 2H,I). Despite having a significantly lower carrying capacity in the field, p95 exhibits an equivalent number of screen-detectable tumors.

The invasive potential of a breast cancer clone can sometimes be established before screen detection^15–17^ resulting in early dissemination and metastasis prior to detection^18–21^. Whole lung imaging of Crainbow cells was performed to quantify the number of Crainbow cells that disseminated to the lung (Figure 2J). Single disseminated cells could be found before palpable tumors could be observed (between 10 and 12 weeks of age) and p95 HER2 expressing cells were 5-fold more numerous in the lung than d16 HER2 (Figure 2J,K). Thus, progression to malignancy is independent of growth, and for p95 may occur abruptly prior to screen detection.

We reasoned that our data could be used to quantify the rates of major transitions during tumor development so that we could quantitatively compare growth models and progression potentials for each genotype. Tumors grow along a continuum from the earliest initiation event, to small, image detectable lesions, and finally to large, palpable tumors (Figure 3A). While we cannot directly trace a single cell or tumor’s growth from start to finish, a statistical population-based model from many distinct points in time can infer tumor transitions from state to state. Based on the assumptions that all small tumors arise from a transformed cell in the field and that all large tumors were once small tumors, a series of ordinary differential equations (ODEs) are defined to capture the surveyed data from Figure 2 and estimate transition rates from one state to the next (Figure 3A,B). Due to the limited number of WT HER2 tumors we once again removed them from this analysis. Based on measurements from Figure 2B, a logistic growth model of the fields of HER2 expressing cells (*F*) was used to describe the exponential growth rate of HER2+ cells (*r)* that occurs during puberty from 4 to 6 weeks of age and then approaches a carrying capacity within the mammary duct (*K*) (Figure 3B,C). d16 HER2 transformed field cells expand at an ∼5 fold higher rate than p95 leading to a 9-fold higher carrying capacity (Figure 3C). The number of small and large tumors over time are plotted by fitting the ODEs to the whole mouse tumor measurements with 90% confidence intervals (Figure 3D,E). d16 screen detectable tumors are more prevalent over time compared to p95 (Figure 3D). In contrast, p95 HER2 large tumor number remains higher than d16 HER2 throughout the observed time frame (Figure 3E).

**Figure 3.**
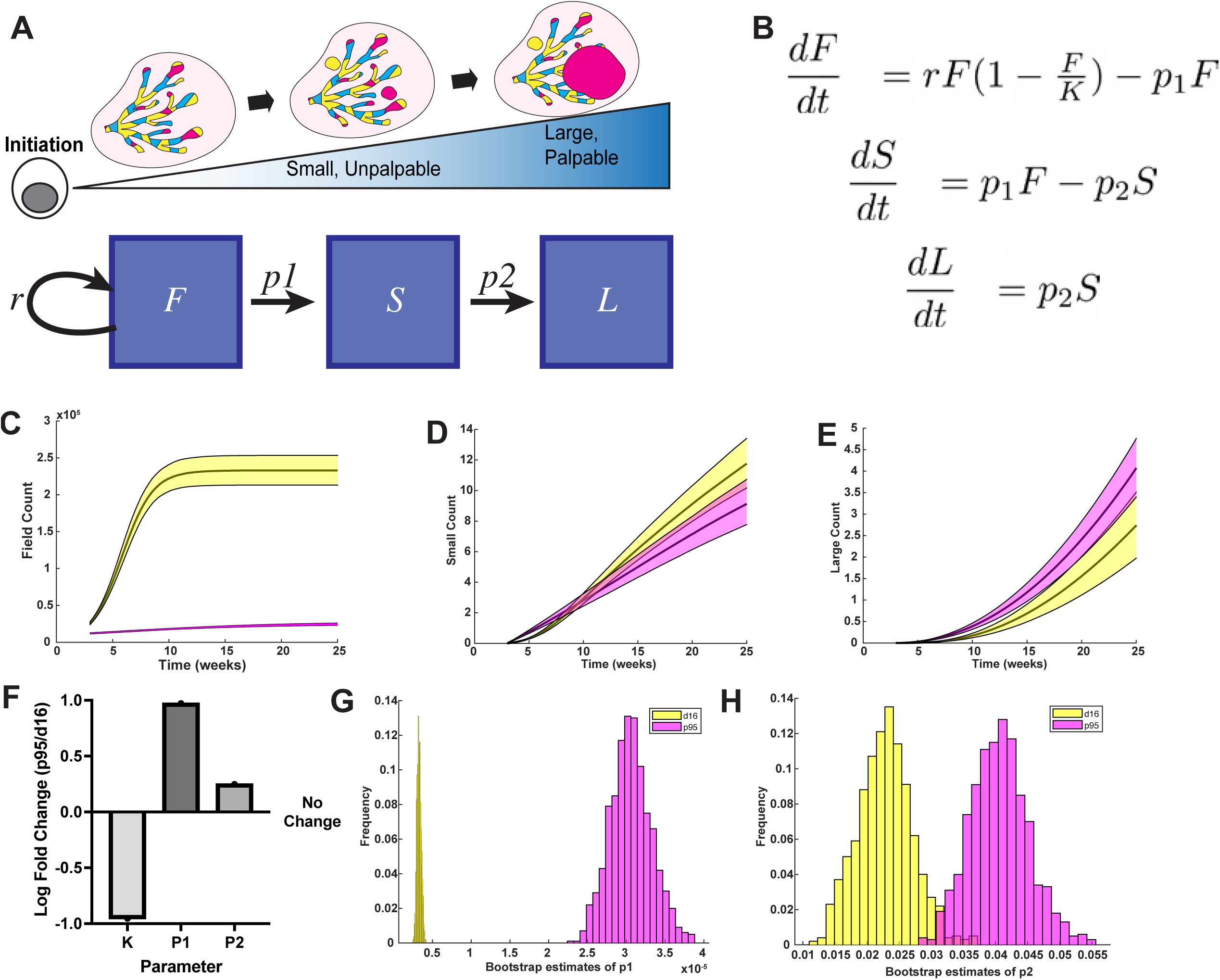
Three compartment model of tumorigenesis. **A,** Schematic of discrete states of tumorigenesis in HBOW mice. Fields of recombined cells (*F)* grow at a rate (*r)* before progressing to small, unpalpable lesions (*S*) at rate *p1*. Small lesions then progress to large, palpable tumors (*L*) at rate, *p2*. **B,** Ordinary differential equations defining progression rates from one compartment to the next. **C,** Estimated logistic growth of fields of cells expressing d16 (yellow) or p95 (magenta) in HBOW mice over time. **D,E** Logarithmic fit of the number of S (screen detectable) (**D**) and L (palpable) lesions (**E**) per mouse across time. **F,** Fold difference between p95 HER2 and d16 HER2 in field size, p1 rate, and p2 rate. **G-H,** Bootstrap estimates of p1 (**G**) and p2 (**H**). **C-E:** Error bars represent 90% confidence intervals.

Our model was then used to determine which state transition described the high p95 tumor burden despite the lack of a large field. Bootstrapping was used to make statistical inferences and estimated progression rates. The progression rate (*p1*) from transformed cell (*F*) to screen detectable tumor (*S*) is a log order higher for p95 compared to d16 HER2 (Figure 3F,G; Table1). Progression rate (*p2*) from small (*S*) to large tumors (*L*) is also two-fold higher than for p95 HER2 (Figure 3F,H; Table 1). Comparison of the progression rates between compartments demonstrates how each successive state transition becomes more likely. The model demonstrates that progression from a transformed field cell to a screen detectable tumor is the rate limiting step in tumorigenesis and that this transition is much more likely for p95 compared to d16 transformed cells.

**Table 1.** Three compartment model parameters with 95% confidence interval.

|  | d16 | CI 95% | p95 | CI 95% |
| --- | --- | --- | --- | --- |
| F0 | 26000 | N/A | 12000 | N/A |
| K | 233000 | N/A | 26000 | N/A |
| r | 0.7313 | N/A | 0.1346 | N/A |
| p1* | 0.000003259 | 2.786e-6, 3.846e-6 | 0.0003048 | 2.544e-5, 3.599e-5 |
| p2* | 0.0229 | 0.0148, 0.0315 | 0.0405 | 0.0321, 0.0502 |

\* denotes significance

Our data suggest that an ill-defined state prior to screen detection may be a critical transition that a transformed cell encounters. For instance, two putative neoplastic states in the mouse mammary gland have been established: hyperplastic alveolar nodules(HAN) and intraductal lesions^22–25^, both of which are below our sensitivity of detection using whole mouse imaging. Confocal imaging of Crainbow fluorescence in FFPE sections from HBOW identifies extensive side-budding reminiscent of HAN that predominantly expresses d16 HER2 as well as lesions within the duct expressing both d16 and p95 HER2. (Figure 4A). Due to the rarity and labile nature of these lesions it is difficult to quantify and as such we define this state as the “occult”. Using our mathematical model, a new compartment for occult dysplastic lesions (O) was incorporated to develop a four-state model of tumorigenesis (Figure 4B).

**Figure 4.**
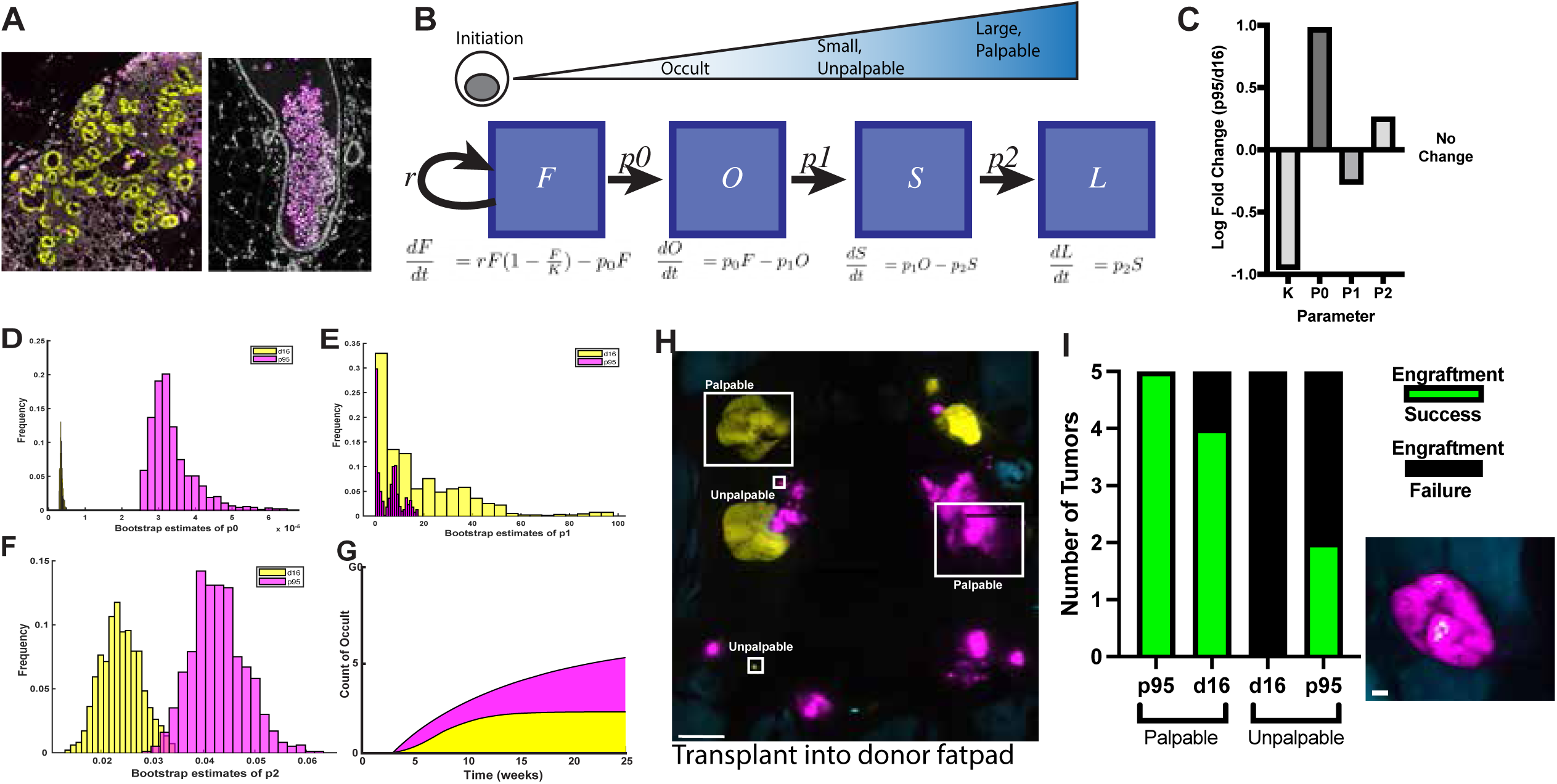
Four compartment model of occult tumorigenesis. **A,** Confocal imaging of potential occult lesions in HBOW mammary glands expressing d16 (yellow) or p95 (magenta). **B,** Schematic of discrete states of tumorigenesis in HBOW mice. Fields of recombined cells (*F)* grow at a rate (*r)* before progressing to occult dysplasia (O) at rate (*p0*). Dysplastic lesions progress to small, unpalpable tumors (*S*) at rate *p1*. Small lesions then progress to large, palpable tumors (*L*) at rate, *p2*. Ordinary differential equations defining progression rates from one compartment to the next are displayed below each compartment. **C,** Fold difference between p95 HER2 and d16 HER2 in field size, p0 rate, p1 rate, and p2 rate.**D-F,** Bootstrap estimates of p0 (**D**), p1 (**E**) and p2 (**F**). **G,** Estimates of the number of occult dysplastic lesions for d16 HER2 and p95 HER2. Error bars represent 80% confidence intervals. **H,** Whole mouse image with representative palpable and unpalpable tumors in boxes. Tumors were removed for digestion and transplantation into donor fatpads. Scale bar = 10mm. **I,** Engraftment rate for tumor cells dissociated from unpalpable and palpable tumors and transplanted into donor mammary fat pads. Green denotes a successful engraftment and black a transplant that failed to grow (n=5). Lower right shows a representative outgrowth from a palpable p95 tumor transplant (magenta). Scale bar = 1mm.

The four-state model revealed that transition out of the field compartment, *p0,* is roughly an order of magnitude higher for p95 than d16, and the transition into the palpable compartment, *p2,* is roughly twice as high for p95 compared to d16 (Figure 4C,D,F; Table 2). Not present in the FSL model, the transition rate from an occult dysplasia to a screen detected neoplasia, *p1,* is about two-fold greater for d16 than p95, though this is not significantly different (Figure 4D,E; Table 2). We used this model to estimate the number of occult (“O”) lesions over time. Estimates of the number of occult lesions are not predictive but suggest that p95 more frequently progresses from the field (F) and generates more occult lesions that readily advance to screen-detectable lesions (Figure 4G). This occurs despite a relatively small reservoir of field cells. Despite the uncertainty in the rate *p1* and the state O, inclusion of an occult compartment highlights that the transition from a field cell to a detectable tumor is not a linear process and each genotype encounters a critical rate limiting step in the tumorigenic process that occurs long before becoming detectable.

**Table 2.** Four compartment model parameters with 95% confidence intervals.

|  |  | d16 | 95% CI | p95 | 95% CI |
| --- | --- | --- | --- | --- | --- |
|  | F0 | 26000 | N/A | 12000 | N/A |
|  | K | 233000 | N/A | 26000 | N/A |
|  | r | 0.7313 | N/A | 0.1346 | N/A |
| C=9mm | p0* | 3.40E-06 | 2.870e-6, 4.339e-6 | 3.20E-05 | 2.652e-5, 4.952e-5 |
|  | p1 | 10.4332 | 0.4220, 65.5811 | 5.6362 | 0.2117, 15.5382 |
|  | p2* | 0.0237 | 0.0178, 0.0310 | 0.0426 | 0.0348, 0.0522 |
| C=1.5mm | p0* | 3.38E-06 | 2.861e-6, 6.940e-6 | 3.21E-05 | 2.632e-5, 5.079e-5 |
|  | p1 | 8.2566 | 0.1101, 99.8215 | 4.7475 | 0.2064, 17.1339 |
|  | p2* | 0.0764 | 0.0614, 0.0981 | 0.1258 | 0.1051, 0.1586 |

\* denotes significance

Next, we determined whether progression to malignancy was associated with transition through the occult compartment. One measure of progression is a test of autonomous growth that can be modeled by transplantation into a syngeneic host^26,27^. Whole mouse imaging was performed to identify paired palpable and unpalpable tumors of the same genotype from the same mouse (N=5 per genotype). These tumors were then dissected and transplanted into the orthotopic fatpad of immune intact and syngeneic FVB/N female mice between 4 and 8 weeks of age (Figure 4H). Successful engraftment was determined by a tumor growing to humane endpoint and maintaining HER2 expression as confirmed by whole mouse imaging. Evaluation of engraftment rate revealed that all palpable p95 tumors successfully engrafted while 80% of d16 tumors transplanted successfully (Figure 4I) suggesting again that at clinical diagnosis both d16 and p95 tumors have progressed to an autonomous state. 40% of p95 unpalpable lesions transplanted successfully while none of d16 unpalpable lesions could be transplanted (Figure 4I). This suggests that tumor progression can occur synchronously with the occult transition resulting in nascent lethal lesions at screen detection.

Lastly, sensitivity analysis was performed in order to understand how variations in parameters (i.e. r, K, p1, p2) affect overall tumorigenic potential ^28,29^. In this analysis, we see the most sensitive parameter for a clinically detectable tumor (*L*) is *p0*, the transition rate from the field to an occult dysplastic lesion, for both the p95 and d16 models (Figure 5A,B). In other words, changes in the occult transition rate have major impacts on tumorigenic potential. An interesting difference between the genotypes is that for d16, the field carrying capacity (*K*) has effects of the same magnitude of the occult transition (*p0*) (Supplementary Figure 2A,B) whereas for p95, *K* and the initial field size (*F0*) are about equal and have half the magnitude than *p0*. This suggests that the development of p95 tumor is less dependent upon carrying capacity (*K*) and the initial field size (*F0*) compared to d16.

**Figure 5.**
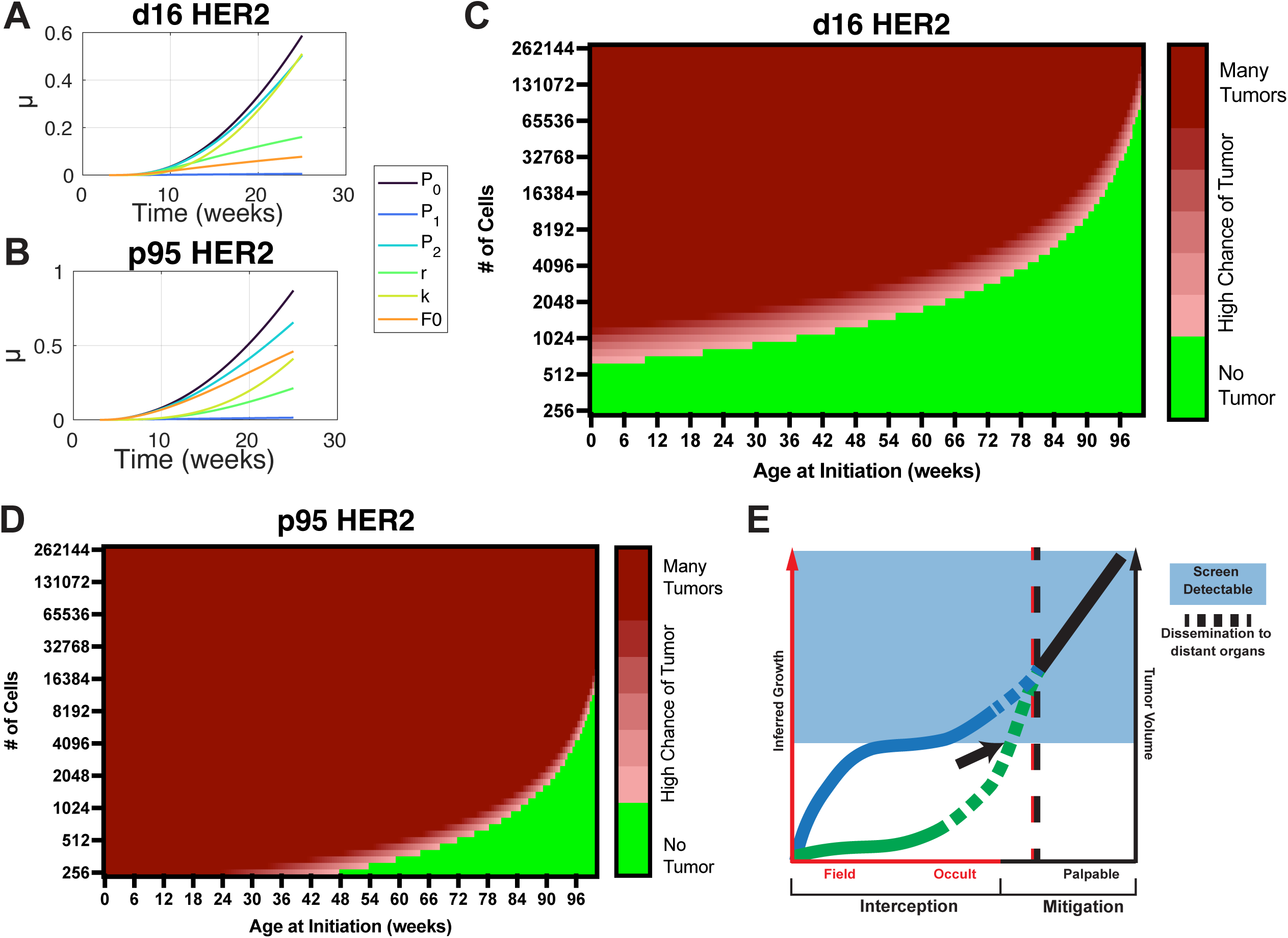
Clinical implications of Crainbow modeling. **A&B,** Sensitivity plots of model fit showing the relative contribution of each variable to the uncertainty in determining transition to palpable tumors expressing d16(**A**) and p95(**B**). **C&D,** Simulated data using the FSL model to visualize the impact of changing both the time of initiation and field size on the likelihood of a tumor forming within the lifetime of a mouse. The simulated data is based on oncogenic cells expressing either d16 (**C**) or p95 (**D).** Green represents a low likelihood of any tumor forming (<0.5). The red gradient represents an increasing likelihood of a tumor developing (0.5-1). Dark red represents 1 or more tumors. **E,** Graphical representation of 2 cases of non-linear tumor progression. Both cases occur prior to the continuous, clinically observable growth of large tumors. The blue line represents a rapidly growing field and its progression to a small screen-detectable tumor. In this case, cancer mitigation by screen detection (blue shade) can occur before dissemination to distant organs (dotted line) and reduce the risk of metastasis. In contrast, the green line represents the slow growing field that rapidly progresses to an invasive lesion that disseminates (dotted line) before becoming screen detectable (Black arrow). This nascent lethal phenotype reduces the efficacy of Mitigation. Thus, cancer interception strategies must be pursued to prevent nascent lethal progression. interception methods, which if able to predict when and where cancer progression will occur, can treat before the transition to a screen detectable lesion and may be more successful than traditional mitigation^31,36^.

We can visualize differences in occult transitions by leveraging the statistical models for d16 or p95 tumor development to simulate the number of palpable tumors that would form from a given field of cells within an average lifetime of a captive mouse (2 years). If the initial event occurs early (∼3 weeks of age), many detectable p95 tumor are likely to develop from a very small field of cells (<256) compared to d16 HER2 which requires nearly 1000 cells to have a high probability of one tumor (Figure 5C,D). If the simulation begins at 1 year of age, in order to have a high likelihood of one tumor, d16 HER2 requires a nearly 6 times larger field of oncogenic cells compared p95 HER2. This difference in number of cells needed to likely form a tumor is representative of the relative frequency at which any oncogenic cell is able to navigate through the relatively difficult occult transition to eventually become a large, invasive tumor. This projection of our data demonstrates how nonlinear progression to an invasive tumor during the occult transition can lead to nascent lethal cancers that confound current cancer screening paradigms.

## Discussion

The benefits and risks of screening are under increasing scrutiny, and in the breast, there is evidence to suggest net neutrality in survival despite decades of intense screening^35^. Three main phases of cancer intervention have recently been summarized and include: prevention, interception, and mitigation^30,31^. The conventional linear progression paradigm supports the strategy of mitigation, using surveillance to detect anatomically small tumors with the expectation that clinically detecting and treating these smaller tumors results in decreased mortality^32–34^. Unfortunately, even after decades of screening for clinically detectable tumors, this strategy has not resulted in the expected positive benefits of reducing mortality or late-stage disease^2^.

Our data demonstrates the paradox of cancer screening using a genetically tractable mouse model system (Figure 5E). We show that tumors can originate from rapidly growing premalignant fields (i.e. d16-HER2). Larger field sizes increase the likelihood that a rare clone advances to a small image detectable event in the mouse that is unlikely to disseminate but if left untreated may eventually metastasize. Therefore, early screening and appropriate interventions could reduce the likelihood of a malignant transition and prevent recurrent, late-stage disease. On the other hand, our modeling also shows how quiescent clones (i.e. p95-HER2) are less likely to rapidly expand. Yet, their ability to transition from premalignancy to malignancy is high. Thus, these nascent lethal clones have already disseminated and are much more likely to escape, prior to anatomic disruption sufficient for clinical detection. In this case,

We also provide experimental evidence for a model of “Biological predeterminism”^37,38^, whereby the lengthy preclinical process establishes lethality prior to detection. Curiously, the most detectable lesions in our model are associated with an isoform of HER2 that is also predictive of favorable therapeutic responses^39^. Thus, d16 driven tumors are likely to have favorable outcomes regardless of early detection. In contrast, a tumor lacking a treatable domain, like p95 HER2^40–42^, is also the most unlikely to be detected, and most likely to disseminate. Thus, in our model, p95 represents the worst-case scenario for a patient – a treatment insensitive tumor that is not screen detectable. This continues to support the notion that p95 is associated with worse clinical outcomes^40,41,43^ and further warrants the need to clinically validate this isoform as a predictive biomarker.

Our model points to an important occult transition where a transformed cell establishes a neoplastic lesion. We chose the term occult because of the inability to image the exact moment of this transition. This leads to a challenging question, what is the occult transition? The occult transition could represent a location in the form of a privileged site of growth. We speculate that an occult compartment could reside inside the duct, similar to the ductal hyperplasias described several decades ago^22^, in which growth inside the duct provides for a unique microenvironment compared to growth into the surrounding fat. For instance, mammary ducts are surrounded by a layer of ductal macrophages^44^. This could provide a mechanism whereby disseminating cells could be in close proximity to macrophage populations necessary for early invasion^45,46^. The occult transition could also represent a moment in time where malignancy is established by changes in the host that impact fitness of tumor cell phenotypes. These could be systemic in nature and may range from immune ageing, microbiome changes, obesity, injury and many others^5^. The occult transition could also occur through oncogene driven differences in cell state plasticity. We also have evidence for early state changes in the p95 expressing mammary epithelium that results in EMT-like states and an early invasive histological phenotype^12^. This suggests that functionally selective signaling across HER2 isoforms may also be important^47^. The challenge of defining the occult transition is key to improving the ability to predict and detect this critical transition.

There are limitations and assumptions to our models that must also be considered. A significant assumption is the cutoff between a screen detectable and palpable tumor in imaging based data. While somewhat experimentally determined from palpation data, we tested this cutoff by utilizing a previously described, anatomically based 1.5mm^2^ cutoff^12^. When applied to our four compartment model, parameter estimates changed by less than 10% and overall tumor transition dynamics were unchanged due to the high impact of the p0 transition in relation to the p2 transition (Supplementary Figure 1). Secondly, our system is an abstraction of a much more complicated tumor ecosystem. Our mouse model overexpresses human HER2 isoforms. Although this is a useful model system of HER2 amplification, it is still not naturalistic. Similarly, the system of ODEs is a mathematical abstraction of the complex tumor ecosystem and has several key limitations, some of which would be common to any similar mathematical model. Primarily, spatial relationships cannot be described by the current ODEs. Additionally, the parameters of the dynamical system are estimated using survey data (mouse necropsy) instead of longitudinal data. Thus, the use of certain statistical frameworks, e.g. nonlinear mixed-effects modeling, that describe the variation between individuals are not applicable^48,49^. Lastly, the models describe observations of 2-dimensional tumor size (mm^2^), not the biological state (e.g. metastatic or benign) or interactions between tumor genotypes. Addressing these limitations to more deeply understand these transitions critical to tumor development are paramount to improving the ability to intercept nascent lethal lesions.

Intercepting the progression of a premalignant clone is a significant challenge but could provide a transformative option for preventing cancer. Overall, we demonstrate the importance of reevaluating major conceptual advances of the past with modern era technological advances. Our work affirms the rules of progression yet provides an experimental framework for future work for predicting progression risk in a patient’s lifetime.

**Supplementary Figure 1.**
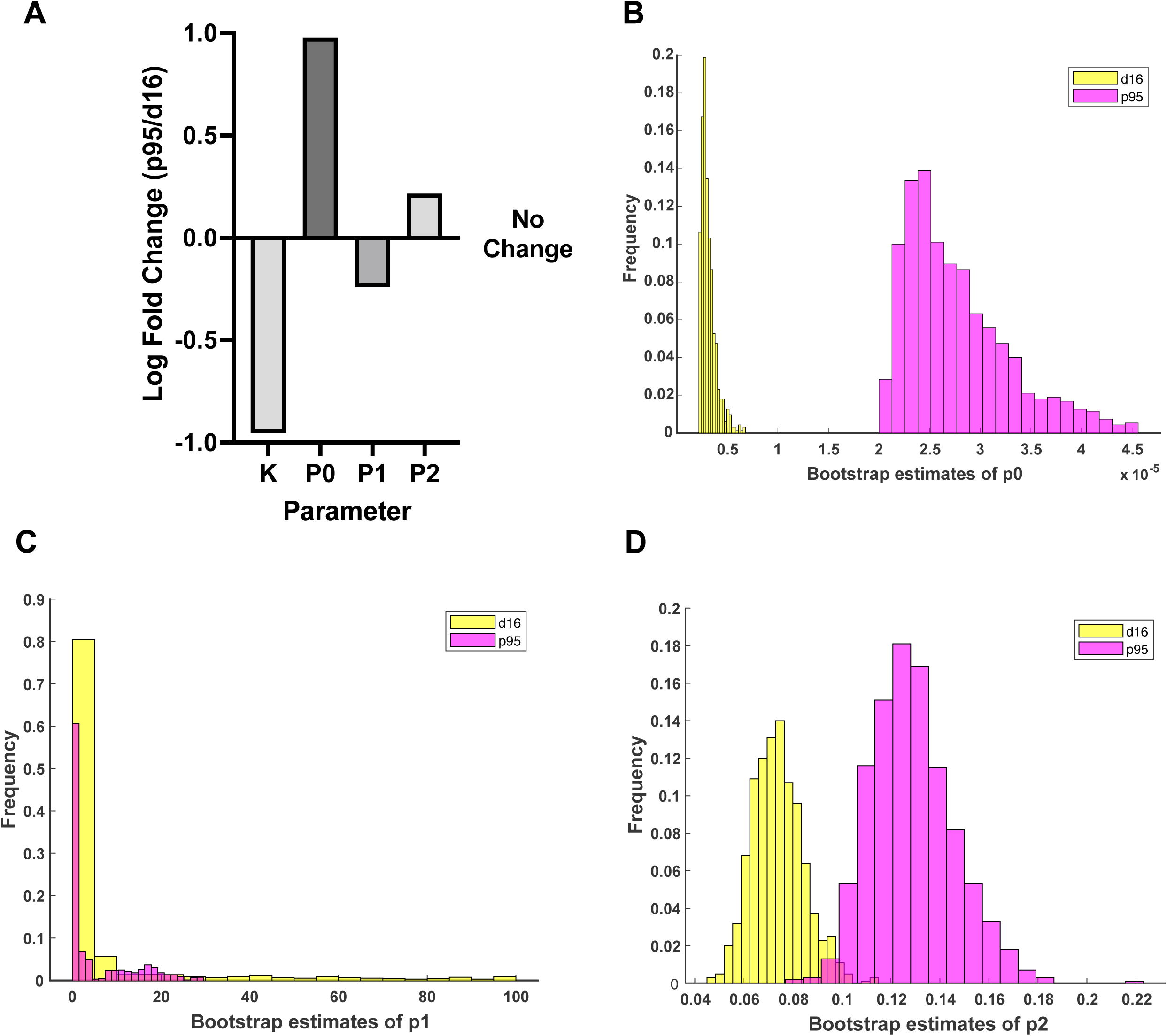
Effect of S:L cutoff in four compartment model of tumorigenesis. **A,** Fold change difference in parameter estimates between p95 and d16 when the cutoff from S:L is 1.5mm^2^. **B-D,** Bootstrap estimates of *p0*(**B**), *p1*(**C**), and *p2*(**D**).

## Materials and Methods

### Animal Studies

All animal studies were performed according to an approved protocol from the Duke Institutional Animal Care and Use Committee. HER2 Crainbow mice were crossed to MMTV Cre (MMTV-cre/Line D (MGI, catalog no. 3581599, RRID:MGI:3581599) to generate HBOW ^12^. Both lines were fully backcrossed to the FVB/N background. All experimental animals were pre-emptively enucleated at 3 weeks of age due to off target harderian gland tumors ^50^. All experimental animals used were female.

### Mammary Gland Epithelial Cell Analysis

Digestion of mammary glands for analysis of epithelial cell number was performed following the protocol from Ludwik et al, 2021^51^. Briefly, all 10 mammary fat pads were removed from M-HER2 mice, finely minced, and incubated in Collagenase A(2mg/mL) in RPMI media for 3 hours at 37C with 5% CO2. Next, digested tissue was incubated in DNase I followed by RBC lysis. Then digested tissue was spun down and resuspended in Accumax for 10 minutes on a rotating shaker at 37C. Next, cells were pelleted and resuspended in 5X Trypsin for 5 minutes at 37C before being filtered through a 40um filter. Samples of single cell suspensions were imaged on a Zeiss 880 confocal and counted using Imaris(v10.1).

### Whole Mouse Imaging

Mice were euthanized with CO2 and pinned on vinyl dissecting pads to expose all 10 mammary glands. Each mouse was then imaged on a Zeiss Axiozoom. Images were imported into either QuPath^52^ or ImageJ2 ^53^ for manual measurement and classification of all tumors.

### Whole Organ Imaging

Mammary glands and lungs were dissected and placed in or inflated with 10% neutral buffered formalin (NBF) for fixation. After 2 hours of fixation, tissues were briefly washed with 0.5% Tween with PBS for 30 minutes before being submerged in FUNGI clearing solution for at least 4 hours ^54^. After clearing, tissues were sandwiched between a glass slide and coverslip for imaging on a Zeiss 880 confocal. For whole lung imaging, spectra were obtained from lambda mode imaging for each Crainbow fluorescent protein and stored as “online fingerprints” for imaging. This process ensures only Crainbow fluorescence was captured and reduces background fluorescence. Imaging of occult lesions was done on FFPE sections of mammary glands fixed in NBF. FFPE sections were deparaffinized and mounted with DAPI nuclear stain before imaging on a Zeiss 880 confocal microscope. Crainbow fluorescence is imageable following FFPE processing with no further amplification or staining ^12^. Analysis was performed using ImageJ2 ^53^ and Imaris (v10.1).

### Mathematical Modeling

#### Sensitivity Analysis

Sensitivity analysis is an analytical technique that explains the uncertainty in the model output as a function of the uncertainty in the model parameters. Sensitivity analysis provides insight into the relationship between the parameters (*r, K, p1, p2*) and the values of the states (F, S, and L). The more sensitive a parameter is, the more effect it has on the state value. If a parameter is *insensitive,* large changes in its value result in small changes to the state values. Here the Morris method is employed to screen the parameters of the system, using the SAFE software package in MATLAB (v9.14; 2023a) ^55^. Morris screening varies one parameter at a time while the others remain constant, and an *elementary effect* is calculated. The elementary effect is the absolute difference in the change in state, divided by the change in the parameter value. The elementary effect is computed at each point in time, and the result is plotted.

#### Parameter Estimation

Parameters for the logistic equation describing the field (r, K, and the initial conditions F(0)) are estimated <u>first</u> using mammary gland digestion data (Figure 2B,3C). Ordinary least squares (OLS) regression is used to fit the solution of the logistic equation to the week 3, 4 and 6 data. The MATLAB function lsqcurvefit is used to find the values *r, K,* and *F*(0) that minimize the least-squared error between the data points and the solution of the logistic equation. For each of the estimated parameters, we must set a range of allowable values, and the ranges are chosen to be roughly the same as the observed data. The time coordinates are shifted so that the initial condition *F*(0) corresponds to an age of 3 weeks, the time of the earliest digestion studies. We assume that growth of the mammary duct is essentially complete by 6 weeks of age, so we choose the upper and lower bounds of *K* according to the range of data at 6 weeks. After deriving estimates of the logistic growth parameters of the field, these values are set and parameter estimates of the transition rates in the S and L compartments, p1 and p2, are calculated (and p0 in the FOSL model). The state transition rates are also estimated by OLS using the MATLAB function lsqcurvefit. Here, we solve the ODEs (Fig 3B,4C) using MATLAB’s ode15s function and then minimize the sum-of-squared differences with the mouse survey data. Every mouse produces 4 data points: (S,L) for both p95 and d16, and the parameters for the p95 and d16 models are fit independently. To produce confidence intervals for the estimations, we employ a hierarchical bootstrap method. There are 120 mice in the survey data set, and we sample N = 120 mice with replacement. Then from each sampled mouse, we re-sample the tumors in that mouse, recording the area of each resampled tumor. The re-sampled tumors of this synthetic mouse are then stratified by genotype and state (S/L) so that each sampled mouse produces 4 data points. The hierarchical bootstrap is repeated 1000 times. Parameters are estimated using median values, and confidence intervals are given by percentiles of the bootstrapped distribution.

To account for additional uncertainty, the field parameters *r,* K, and F(0) are resampled at each bootstrap iteration, using a normal distribution with mean given by the preliminary OLS estimate and standard deviation equal to 5% of the mean. Uncertainty in the mouse age is modeled by adding a normally distributed random number with mean 0 and std. 0.05 (weeks) to each synthetic mouse’s age at every bootstrap iteration.

#### Simulations

To produce the results in Figures 5 C and D, we use the parameter estimates for p1 and p2 in the FSL model and forward simulate the system from the Age at initiation, given on the x-axis, to 2 years, the expected lifespan of a mouse. Here, K is set to the observed number of transformed cells (y-axis) so that the growth of the field (F) is not considered as part of the simulation results.

### Transplant Studies

Tumors to be used for transplant were selected from Axiozoom imaging and digested following the same digestion protocol as described in **Mammary Gland Epithelial Cell Analysis**. Single cells were resuspended in USP saline. Female FVBN/J mice (Jackson Labs) between 4 and 8 weeks of age were anesthetized using Isoflurane and 500,000 isolated tumor cells were injected into the right, fourth(inguinal) mammary gland. Tumors were measured once weekly. At tumor burden endpoint, **Whole Mouse Imaging** was performed to assess genotype.

## Resource Availability

Any requests for resources or data should be directed to and will be fulfilled by Joshua Snyder. The HER2 Crainbow mouse model is available upon request and with a completed materials transfer agreement. This study did not generate any new unique reagents. Data and code will be made publicly available upon publication.

## Acknowledgements

Funding for this work was provided by NCI 1R01CA255372-01 to JCS and DoD W81XWH-21-2-0031to HKL.

## Author contributions

Conceptualization, JDG, LSB, HKL, BWR, and JCS; Methodology, JDG, LSB, BWR, and JCS; Software, HC and BWR; Validation, JDG, HC, and BWR. Formal Analysis, JDG, HC, BWR, and JCS; Investigation, JDG and EJA; Resources, JES, HKL, BWR, and JCS; Data Curation, JDG, HC, and BWR; Writing – Original Draft, JDG, BWR, and JCS; Writing – Review & Editing, JDG, ADB, RDC, HKL, and JCS; Visualization, JDG, HC, BWR, and JCS; Supervision, JDG, BWR, and JCS; Project Administration, JCS. Funding Acquisition, HKL and JCS.

## References

1. Gil Del Alcazar, C.R., Huh, S.J., Ekram, M.B., Trinh, A., Liu, L.L., Beca, F., Zi, X., Kwak, M., Bergholtz, H., Su, Y., et al. (2017). Immune Escape in Breast Cancer During In Situ to Invasive Carcinoma Transition. Cancer Discovery 7, 1098–1115. 10.1158/2159-8290.CD-17-0222.

2. Welch, H.G., Kramer, B.S., and Black, W.C. (2019). Epidemiologic Signatures in Cancer. N Engl J Med 381, 1378–1386. 10.1056/NEJMsr1905447.

3. Welch, H.G., Prorok, P.C., O’Malley, A.J., and Kramer, B.S. (2016). Breast-Cancer Tumor Size, Overdiagnosis, and Mammography Screening Effectiveness. N Engl J Med 375, 1438–1447. 10.1056/NEJMoa1600249.

4. Gerstung, M., Jolly, C., Leshchiner, I., Dentro, S.C., Gonzalez, S., Rosebrock, D., Mitchell, T.J., Rubanova, Y., Anur, P., Yu, K., et al. (2020). The evolutionary history of 2,658 cancers. Nature 578, 122–128. 10.1038/s41586-019-1907-7.

5. Swanton, C., Bernard, E., Abbosh, C., André, F., Auwerx, J., Balmain, A., Bar-Sagi, D., Bernards, R., Bullman, S., DeGregori, J., et al. (2024). Embracing cancer complexity: Hallmarks of systemic disease. Cell 187, 1589–1616. 10.1016/j.cell.2024.02.009.

6. Huels, D.J., Bruens, L., Hodder, M.C., Cammareri, P., Campbell, A.D., Ridgway, R.A., Gay, D.M., Solar-Abboud, M., Faller, W.J., Nixon, C., et al. (2018). Wnt ligands influence tumour initiation by controlling the number of intestinal stem cells. Nat Commun 9, 1132. 10.1038/s41467-018-03426-2.

7. van Neerven, S.M., de Groot, N.E., Nijman, L.E., Scicluna, B.P., van Driel, M.S., Lecca, M.C., Warmerdam, D.O., Kakkar, V., Moreno, L.F., Vieira Braga, F.A., et al. (2021). Apc-mutant cells act as supercompetitors in intestinal tumour initiation. Nature. 10.1038/s41586-021-03558-4.

8. Brown, S., Pineda, C.M., Xin, T., Boucher, J., Suozzi, K.C., Park, S., Matte-Martone, C., Gonzalez, D.G., Rytlewski, J., Beronja, S., et al. (2017). Correction of aberrant growth preserves tissue homeostasis. Nature 548, 334–337. 10.1038/nature23304.

9. Hill, W., Lim, E.L., Weeden, C.E., Lee, C., Augustine, M., Chen, K., Kuan, F.-C., Marongiu, F., Evans, E.J., Moore, D.A., et al. (2023). Lung adenocarcinoma promotion by air pollutants. Nature 616, 159–167. 10.1038/s41586-023-05874-3.

10. Evans, E.J., and DeGregori, J. (2021). Cells with Cancer-associated Mutations Overtake Our Tissues as We Age. Aging Cancer 2, 82–97. 10.1002/aac2.12037.

11. Boone, P.G., Rochelle, L.K., Ginzel, J.D., Lubkov, V., Roberts, W.L., Nicholls, P.J., Bock, C., Flowers, M.L., von Furstenberg, R.J., Stripp, B.R., et al. (2019). A cancer rainbow mouse for visualizing the functional genomics of oncogenic clonal expansion. Nat Commun 10, 5490. 10.1038/s41467-019-13330-y.

12. Ginzel, J.D., Acharya, C.R., Lubkov, V., Mori, H., Boone, P.G., Rochelle, L.K., Roberts, W.L., Everitt, J.I., Hartman, Z.C., Crosby, E.J., et al. (2021). HER2 Isoforms Uniquely Program Intratumor Heterogeneity and Predetermine Breast Cancer Trajectories During the Occult Tumorigenic Phase. Mol Cancer Res. 10.1158/1541-7786.MCR-21-0215.

13. Förnvik, D., Lång, K., Andersson, I., Dustler, M., Borgquist, S., and Timberg, P. (2016). ESTIMATES OF BREAST CANCER GROWTH RATE FROM MAMMOGRAMS AND ITS RELATION TO TUMOUR CHARACTERISTICS. Radiat Prot Dosimetry 169, 151–157. 10.1093/rpd/ncv417.

14. Kay, K., Dolcy, K., Bies, R., and Shah, D.K. (2019). Estimation of Solid Tumor Doubling Times from Progression-Free Survival Plots Using a Novel Statistical Approach. AAPS J 21, 27. 10.1208/s12248-019-0302-5.

15. Casasent, A.K., Schalck, A., Gao, R., Sei, E., Long, A., Pangburn, W., Casasent, T., Meric-Bernstam, F., Edgerton, M.E., and Navin, N.E. (2018). Multiclonal Invasion in Breast Tumors Identified by Topographic Single Cell Sequencing. Cell 172, 205–217.e12. 10.1016/j.cell.2017.12.007.

16. Wang, K., Kumar, T., Wang, J., Minussi, D.C., Sei, E., Li, J., Tran, T.M., Thennavan, A., Hu, M., Casasent, A.K., et al. (2023). Archival single-cell genomics reveals persistent subclones during DCIS progression. Cell 186, 3968–3982.e15. 10.1016/j.cell.2023.07.024.

17. Lips, E.H., Kumar, T., Megalios, A., Visser, L.L., Sheinman, M., Fortunato, A., Shah, V., Hoogstraat, M., Sei, E., Mallo, D., et al. (2022). Genomic analysis defines clonal relationships of ductal carcinoma in situ and recurrent invasive breast cancer. Nat Genet 54, 850–860. 10.1038/s41588-022-01082-3.

18. Hu, Z., Li, Z., Ma, Z., and Curtis, C. (2020). Multi-cancer analysis of clonality and the timing of systemic spread in paired primary tumors and metastases. Nat Genet 52, 701–708. 10.1038/s41588-020-0628-z.

19. Klein, C.A. (2020). Cancer progression and the invisible phase of metastatic colonization. Nat Rev Cancer 20, 681–694. 10.1038/s41568-020-00300-6.

20. Klein, C.A. (2009). Parallel progression of primary tumours and metastases. Nat Rev Cancer 9, 302–312. 10.1038/nrc2627.

21. Harper, K.L., Sosa, M.S., Entenberg, D., Hosseini, H., Cheung, J.F., Nobre, R., Avivar-Valderas, A., Nagi, C., Girnius, N., Davis, R.J., et al. (2016). Mechanism of early dissemination and metastasis in Her2+ mammary cancer. Nature 540, 588–592. 10.1038/nature20609.

22. Medina, D. (2000). The Preneoplastic Phenotype in Murine Mammary Tumorigenesis. J Mammary Gland Biol Neoplasia 5, 393–407. 10.1023/A:1009529928422.

23. Cardiff, R.D., and Wellings, S.R. (1999). The Comparative Pathology of Human and Mouse Mammary Glands. Journal of Mammary Gland Biology and Neoplasia 4, 105–122. 10.1023/a:1018712905244.

24. Cardiff, R.D., Wellings, S.R., and Faulkin, L.J. (1977). Biology of breast preneoplasia. Cancer 39, 2734–2746. 10.1002/1097-0142(197706)39:6<2734::AID-CNCR2820390661>3.0.CO;2-U.

25. Serial Transplantation of Chemical Carcinogen-Induced Mouse Mammary Ductal Dysplasias2 (1979). JNCI: Journal of the National Cancer Institute. 10.1093/jnci/62.2.397.

26. Borowsky, A.D. (2011). Choosing a Mouse Model: Experimental Biology in Context--The Utility and Limitations of Mouse Models of Breast Cancer. Cold Spring Harbor Perspectives in Biology 3, a009670–a009670. 10.1101/cshperspect.a009670.

27. Horava, A., and Skoryna, S.C. (1955). Observations on the Pathogenesis of Neoplasia. Canadian Medical Association Journal 73, 630.

28. Dickinson, R.P., and Gelinas, R.J. (1976). Sensitivity analysis of ordinary differential equation systems—A direct method. Journal of Computational Physics 21, 123–143. 10.1016/0021-9991(76)90007-3.

29. Morris, M.D. (1991). Factorial Sampling Plans for Preliminary Computational Experiments. Technometrics 33, 161–174. 10.1080/00401706.1991.10484804.

30. Castle, P.E., Faupel-Badger, J.M., Umar, A., and Rebbeck, T.R. (2024). A Proposed Framework and Lexicon for Cancer Prevention. Cancer Discovery 14, 594–599. 10.1158/2159-8290.CD-23-1492.

31. Domchek, S.M., and Vonderheide, R.H. (2024). Advancing Cancer Interception. Cancer Discovery 14, 600–604. 10.1158/2159-8290.CD-24-0015.

32. Fearon, E.R., and Vogelstein, B. (1990). A genetic model for colorectal tumorigenesis. Cell 61, 759–767. 10.1016/0092-8674(90)90186-i.

33. Armitage, P., and Doll, R. (1954). The age distribution of cancer and a multi-stage theory of carcinogenesis. British journal of cancer 8, 1–12. 10.1038/bjc.1954.1.

34. Nordling, C.O. (1953). A new theory on cancer-inducing mechanism. British journal of cancer 7, 68–72. 10.1038/bjc.1953.8.

35. Welch, H.G., and Dey, T. (2023). Testing Whether Cancer Screening Saves Lives: Implications for Randomized Clinical Trials of Multicancer Screening. JAMA Internal Medicine 183, 1255– 1258. 10.1001/jamainternmed.2023.3781.

36. Castle, P.E., Faupel-Badger, J.M., Umar, A., and Rebbeck, T.R. (2024). A Proposed Framework and Lexicon for Cancer Prevention. Cancer Discovery 14, 594–599. 10.1158/2159-8290.CD-23-1492.

37. Macdonald, I. (1966). The natural history of mammary carcinoma. 111, 435–442. 10.1016/s0002-9610(66)80023-5.

38. Macdonald, I. (1951). Biological predeterminism in human cancer. Surg Gynecol Obstet 92, 443–452.

39. Castagnoli, L., Iezzi, M., Ghedini, G.C., Ciravolo, V., Marzano, G., Lamolinara, A., Zappasodi, R., Gasparini, P., Campiglio, M., Amici, A., et al. (2014). Activated d16HER2 homodimers and SRC kinase mediate optimal efficacy for trastuzumab. Cancer Res 74, 6248–6259. 10.1158/0008-5472.CAN-14-0983.

40. Abraham, J., Montero, A.J., Jankowitz, R.C., Salkeni, M.A., Beumer, J.H., Kiesel, B.F., Piette, F., Adamson, L.M., Nagy, R.J., Lanman, R.B., et al. (2019). Safety and Efficacy of T-DM1 Plus Neratinib in Patients With Metastatic HER2-Positive Breast Cancer: NSABP Foundation Trial FB-10. Journal of clinical oncology : official journal of the American Society of Clinical Oncology 37, 2601–2609. 10.1200/JCO.19.00858.

41. Scaltriti, M., Rojo, F., Ocaña, A., Anido, J., Guzman, M., Cortes, J., Di Cosimo, S., Matias-Guiu, X., Ramon y Cajal, S., Arribas, J., et al. (2007). Expression of p95HER2, a Truncated Form of the HER2 Receptor, and Response to Anti-HER2 Therapies in Breast Cancer. JNCI: Journal of the National Cancer Institute 99, 628–638. 10.1093/jnci/djk134.

42. Pedersen, K., Angelini, P.D., Laos, S., Bach-Faig, A., Cunningham, M.P., Ferrer-Ramon, C., Luque-Garcia, A., Garcia-Castillo, J., Parra-Palau, J.L., Scaltriti, M., et al. (2009). A naturally occurring HER2 carboxy-terminal fragment promotes mammary tumor growth and metastasis. Mol Cell Biol 29, 3319–3331. 10.1128/MCB.01803-08.

43. Carvajal-Hausdorf, D.E., Schalper, K.A., Pusztai, L., Psyrri, A., Kalogeras, K.T., Kotoula, V., Fountzilas, G., and Rimm, D.L. (2015). Measurement of Domain-Specific HER2 (ERBB2) Expression May Classify Benefit From Trastuzumab in Breast Cancer. J Natl Cancer Inst 107. 10.1093/jnci/djv136.

44. Dawson, C.A., Pal, B., Vaillant, F., Gandolfo, L.C., Liu, Z., Bleriot, C., Ginhoux, F., Smyth, G.K., Lindeman, G.J., Mueller, S.N., et al. (2020). Tissue-resident ductal macrophages survey the mammary epithelium and facilitate tissue remodelling. Nat Cell Biol 22, 546–558. 10.1038/s41556-020-0505-0.

45. Linde, N., Casanova-Acebes, M., Sosa, M.S., Mortha, A., Rahman, A., Farias, E., Harper, K., Tardio, E., Reyes Torres, I., Jones, J., et al. (2018). Macrophages orchestrate breast cancer early dissemination and metastasis. Nature Communications 9, 21. 10.1038/s41467-017-02481-5.

46. Sidani, M., Wyckoff, J., Xue, C., Segall, J.E., and Condeelis, J. (2006). Probing the microenvironment of mammary tumors using multiphoton microscopy. J Mammary Gland Biol Neoplasia 11, 151–163. 10.1007/s10911-006-9021-5.

47. Dankort, D., Maslikowski, B., Warner, N., Kanno, N., Kim, H., Wang, Z., Moran, M.F., Oshima, R.G., Cardiff, R.D., and Muller, W.J. (2001). Grb2 and Shc adapter proteins play distinct roles in Neu (ErbB-2)-induced mammary tumorigenesis: implications for human breast cancer. Mol Cell Biol 21, 1540–1551. 10.1128/MCB.21.5.1540-1551.2001.

48. Pinheiro, J.C., and Bates, D.M. (1995). Approximations to the Log-Likelihood Function in the Nonlinear Mixed-Effects Model. Journal of Computational and Graphical Statistics 4, 12–35. 10.2307/1390625.

49. Comets, E., Lavenu, A., and Lavielle, M. (2017). Parameter Estimation in Nonlinear Mixed Effect Models Using saemix, an R Implementation of the SAEM Algorithm. Journal of Statistical Software 80, 1–41. 10.18637/jss.v080.i03.

50. Garner, A., Ginzel, J.D., Snyder, J.C., Everitt, J.I., and Landon, C.D. (2022). Veterinary Management of Harderian Gland Tumors in Cancer Rainbow (crainbow) HER2-Positive Mice. Comparative Medicine 72, 403–409. 10.30802/AALAS-CM-22-000061.

51. Ludwik, K.A., Sandusky, Z.M., Wright, E.B., and Lannigan, D.A. (2021). FACS protocol for direct comparison of cell populations isolated from mice. STAR Protocols 2, 100270. 10.1016/j.xpro.2020.100270.

52. Bankhead, P., Loughrey, M.B., Fernández, J.A., Dombrowski, Y., McArt, D.G., Dunne, P.D., McQuaid, S., Gray, R.T., Murray, L.J., Coleman, H.G., et al. (2017). QuPath: Open source software for digital pathology image analysis. Sci Rep 7, 16878. 10.1038/s41598-017-17204-5.

53. Rueden, C.T., Schindelin, J., Hiner, M.C., DeZonia, B.E., Walter, A.E., Arena, E.T., and Eliceiri, K.W. (2017). ImageJ2: ImageJ for the next generation of scientific image data. BMC Bioinformatics 18, 529. 10.1186/s12859-017-1934-z.

54. Rios, A.C., Capaldo, B.D., Vaillant, F., Pal, B., van Ineveld, R., Dawson, C.A., Chen, Y., Nolan, E., Fu, N.Y., Jackling, F.C., et al. (2019). Intraclonal Plasticity in Mammary Tumors Revealed through Large-Scale Single-Cell Resolution 3D Imaging. Cancer Cell 35, 953. 10.1016/j.ccell.2019.05.011.

55. Pianosi, F., Sarrazin, F., and Wagener, T. (2015). A Matlab toolbox for Global Sensitivity Analysis. Environmental Modelling & Software 70, 80–85. 10.1016/j.envsoft.2015.04.009.

